# The distribution of the number of mutations in the genealogy of a sample from a single population

**DOI:** 10.1101/2025.02.04.636538

**Authors:** Yun-Xin Fu

## Abstract

The number *K* of mutations in the genealogy of a sample of *n* sequences from a single population is one essential summary statistics in molecular population genetics and is equal to the number of segregating sites in the sample under the infinite-sites model. Although its expectation and variance are the most widely utilized properties, its sampling formula (i.e., probability distribution) is the foundation for all explorations related to *K*. Recently, it has been established that *K* is subject to the Central Limit Theorem, and thus has asymptotic normality. However, due to its slow convergence to normality, the finite-sample distribution remains indispensable. Although an analytic sampling formula exists, its numerical application is limited due to susceptibility to error propagation. This paper presents a new sampling formula for *K* in a random sample of DNA sequences from a neutral locus without recombination, taken from a single population evolving according to the Wright-Fisher model with a constant effective population size, or the constant-in-state model, which allows the effective population size to vary across different coalescent states. The new sampling formula is expressed as the sum of the probabilities of the various ways mutations can manifest in the sample genealogy and achieves simplicity by partitioning mutations into hypothetical atomic clusters that cannot be further divided. Under the Wright-Fisher model with a constant effective population size, the new sampling formula is closely analogous to the celebrated Ewens’ sampling formula for the number of distinct alleles in a sample. Numerical computation using the new sampling formula is accurate and is limited only by the burden of enumerating a large number of partitions of a large *K*. However, significant improvement in efficiency can be achieved by prioritizing the enumeration of partitions with a large number of parts.

---

The number *K* of mutations in the genealogy of a sample of *n* sequences from a population is one of the most important summary statistics in molecular population genetics. It is also ubiquitous in the analysis of DNA sequence samples, as it can be identified as the number of segregating sites in a sample under the infinite-sites model (Ewens (2004), and also see Fu (2025) for a concise list of theoretical and methodological studies related to *K*). Therefore, it is essential to characterize its sampling formula (i.e., probability distribution), which is

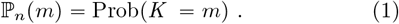

Watterson (1975) established that *K* = *X*_2_ *X*_*n*_, where *X*_*i*_ represents the number of mutations in the time period of observing *i* ancestral sequences in a forward process. In the terminology of the Kingman coalescent (Kingman, 1982a,b), *X*_*i*_ is the number of mutations during state *i* of the coalescent process of the sample, and follows a geometric distribution with parameter 1 − *φ*_*i*_, where

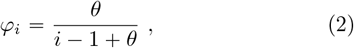

where *θ* = 4*Nµ* with *N* as the effective population size and *µ* as the mutation rate per sequence per generation. Since *X*_*i*_ is independent *X*_*j*_ (*i* ≠ *j*), the probability generating function of *K* is thus the product of those for different *X*_*i*_, which led Watterson (1975) to derive the expectation and variance of *K*, and also pointed out that

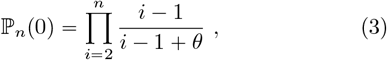

which, as well as for ℙ_*n*_(*m*) for small *m*, can also be obtained from direct convolution. Using partial expansion of the probability generating function of *K*, Tavaré (1984) derived the following sampling formula

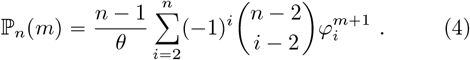

Recently, Fu (2025) established that central limit theory holds for *K*, meaning that *K* asymptotically follows a normal distribution. Furthermore, the finite-sample skewness and kurtosis of *K* are now available, which enhances our understanding of its finite-sample properties. The characterization of *K* appears to be nearly complete, at least under the Wright-Fisher model with a constant effective population size. However, since *K* under the infinite-sites model is often considered a counterpart to *k*, the number of distinct alleles in the sample under the infinite-alleles model, the lack of resemblance of Tavaré’s formula (elegant in its own right) to Ewens’ sampling formula leaves much to be desired.

For a sample configuration 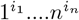 of *n* sequences, where *i*_*j*_ is the number of distinct alleles with *j* copies in the sample, it is known (Ewens, 1972; Karlin and McGregor, 1972) that

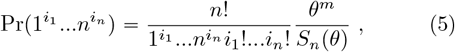

where

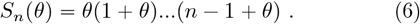

Summing up all sample configurations that have the same number of distinct alleles leads to

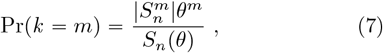

where

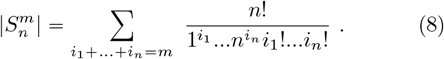

The fact that *K* arises in different configurations, and that its probability is the sum of the probabilities of these configurations, is both intuitive and intriguing. This prompts one to wonder whether a similar analogy exists for *K*. Of course, since *K* is only asymptotically sufficient for *θ* (Fu and Li, 1993), any such sampling formulas, if they exist, are expected to be more complex than the Ewens’ sampling formula.

It is also unfortunate that Tavaré’s sampling formula (4) has a limited scope of applicability in the numerical calculation of ℙ_*n*_(*m*), due to error accumulation in the summation of large absolute values with alternating signs. This issue becomes progressively more severe with increasing sample size, often making the formula unusable even for modest sample sizes. Furthermore, it is highly desirable to be able to compute ℙ_*n*_(*m*) for a non-constant population.

The primary purpose of this paper is to present and prove a new sampling formula for *K*, which not only provides a direct analogy to the Ewens’ sampling formula, but also serves as a reliable method for computing ℙ_*n*_(*m*) under a more general model than the Wright-Fisher model with a constant effective population size.

## The main results

The Kingman coalescent (Kingman, 1982a,b) is best known for describing a sample of size *n* taken from a population that evolves according to the Wright-Fisher model (Ewens, 2004), although the results are robust under several alternative models, including Moran’s model. For this reason, we assume the population evolves according to the *constant-in-state* model (Fu, 2025), which is a variation of the Wright-Fisher model. In this model, the effective population size during coalescent state *i* is constant (*N*_*i*_), while the assumption of independence among coalescent times remains valid. The constant-in-state model extends the Wright-Fisher model by allowing for populations of non-constant sizes, while still maintaining mathematical tractability. As a result, it has been applied in various contexts (Pybus et al., 2000; Liu and Fu, 2015). Throughout the paper, we restrict our attention to a sample of DNA sequences from a single locus without recombination, where all mutations are selectively neutral.

Integer partitions are essential in this paper and thus warrant a brief introduction. A partition of an integer *m* refers to expressing *m* as a sum of positive integers, where the order of the summands does not matter (Riordan, 1958; Abramowitz and Stegun, 1972). For example, 5 = 1 + 2 + 2 = 4 + 1, so the tuples (1, 2, 2) and (4, 1) are both partitions of 5. The order of parts in a partition is irrelevant, and by convention, the parts are listed in descending order. We will use bold letters to represent integer partitions in this paper and adopt the convention that ***s*** ⊢ *m* indicates that ***s*** is a partition of *m*. Thus, ***s*** ⊢ *m* represents a tuple of positive integer(s)

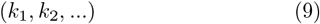

such that *k*_1_ ⩾ *k*_2_ ⩾ … and *k*_1_ = *m*. Equivalent but more convenient notation for an integer partition is cycle type (multiplicity form)

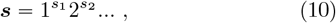

where *s*_*k*_ is the multiplicity (number of occurrences) of integer *k* in the partition tuple. Therefore, the sample configuration 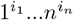 in Eq.(5) is an integer partition of sample size *n*. For example, ***s*** = (5, 2, 2, 1)) 10 has cycle type 1^1^2^2^3^0^4^0^5^1^. It is customary to omit the component 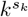 in the cycle type when *s*_*k*_ = 0, so the above cycle type becomes more concisely ***s*** = 1^1^2^2^5^1^. It is also conventional to consider 1^0^2^0^… as the sole partition of integer 0.

Summation (∑) and multiplication (∏) appear frequently in this paper, often in the context of enumerating integer partitions and counting permutations of cycle type (which will be elaborated on later). We adopt the convention that when there is no summand, ∑ is equal to 0, and when there is no factor, ∏ is equal to 1.

The main result of this paper is as follows.

### Theorem.

*Let* ℙ_*n*_(*m*) *denote the probability of having exactly m mutations in the genealogy of a random sample of n sequences from a population evolving according to the constant-in-state model. Then*,

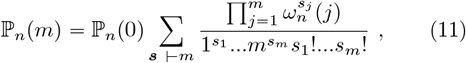

*where the summation is taken over all integer partitions* 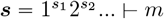 *and*

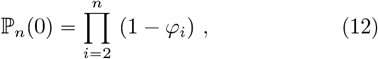

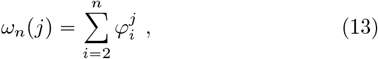

*where*

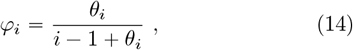

*where θ*_*i*_ = 4*N*_*i*_*µ, with N*_*i*_ *being the effective population size at state i of the coalescent process and µ being the mutation rate per sequence per generation*.

Before proceeding with the proof of the Theorem, we introduce some notations for brevity and clarity. Additionally, we present a corollary for the specific case where the population evolves according to the Wright-Fisher model with a constant effective population size.

Define for 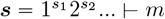

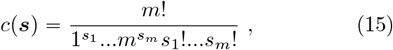

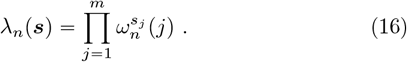

Then Eq.(11) can be written as

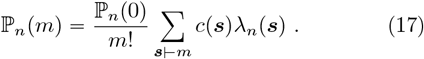

It should be noted that *c*(***s***) defined above is the number of permutations of *m* different objects conforming to cycle type ***s*** (Riordan, 1958; Abramowitz and Stegun, 1972). Its precise meaning in the context of mutations will be elaborated later when we discuss combinatorial interpretation of the Theorem. As far as *c*(***s***) is concerned, there is no need to specify explicitly the integer to which a partition ***s*** belongs, because the integer is equal to

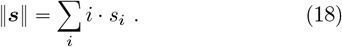

Furthermore it is well known(Abramowitz and Stegun, 1972) that

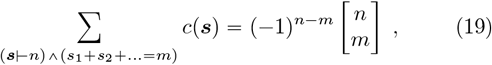

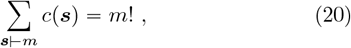

where 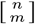 is the Stirling number of the first kind. It follows that 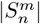 defined by (8) is equal to 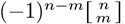, which is known as unsigned Stirling number of the first kind.

Let

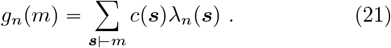

Then enumerating over all integer partitions for a given *m*, it can be verified easily that

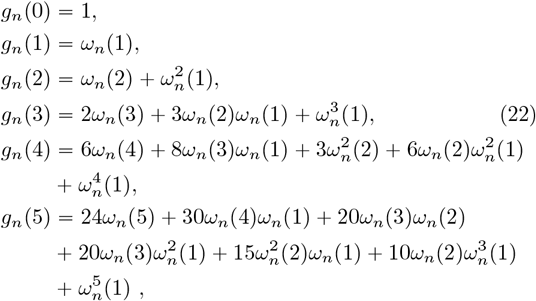

which can be used to compute ℙ_*n*_(*m*) quickly for *m* ⩽5.

Under the Wright-Fisher model with constant effective size, we have that *θ* = *θ*_2_ = … = *θ*_*n*_. It follows from (3) that

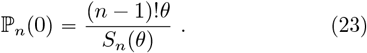

Furthermore all the summands of *ω*_*n*_(*j*) has a common factor *θ*^*j*^ and for a partition ***s***) *m*,

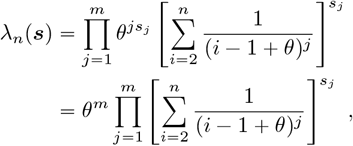

which leads to the following corollary

### Corollary.

*For a random sample of n sequences from a population evolving according to the Wright-Fisher model with constant effective size N, the probability of having exactly m mutations in the sample genealogy is*

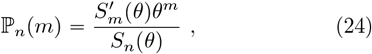

*where S*_*n*_(*θ*) *is defined by (6) and*

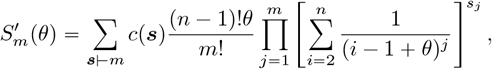

*where the summation is taken over all partitions of m, c*(***s***) *is defined by (15) and θ* = 4*Nµ with µ being the mutation rate per sequence per generation*.

We thus have a closely analogous formula to Ewens’ sampling formula (7), with two key differences. The first is that *k* can take value between 1 and *n*, while *K* can take values between 0 and 8. The second is that, although both 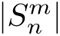 and 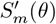 are of the form

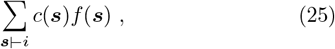

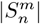 is a summation over partitions of sample size *n* with *m* parts where *f* (***s***) = 1, resulting in the unsigned Stirling number of the second kind (19). In comparison 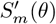 is a summation over all partitions of mutation number *m* with *f* (***s***) as a function of *θ*, partition ***s*** and sample size *n*. This second difference reflects the fact that *k* is a sufficient summary statistic for *θ*, so that the detailed configurations leading to *k* is independent of *θ*, while *K* is not a sufficient summary statistic for *θ*.

### Proof of the Theorem

For clarity, let *K*_*n*_ = *K* for sample size *n*. Consider first the case of *n* = 2. Since *K*_2_ = *X*_2_, which follows a geometric distribution with parameter 1 − *φ*_2_, it follows that

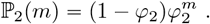

Since 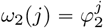,we have for any partition 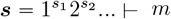 that

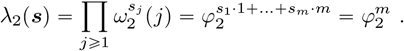

Due to Eq.(20) and *p*_2_(0) = (1 − *φ*_2_), one arrives at

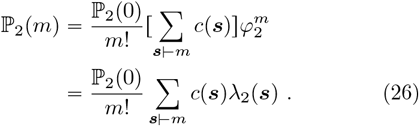

Therefore Eq.(17) is true for *n* = 2. Suppose it is true for all the sample sizes up to *n* − 1. Then since *K*_*n*_ = *K*_*n*− 1_ + *X*_*n*_, the following recurrence relationship holds

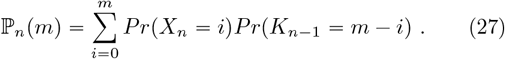

Since *X*_*n*_ ∼ *Geom* (1 − *φ*_2_) it follows that

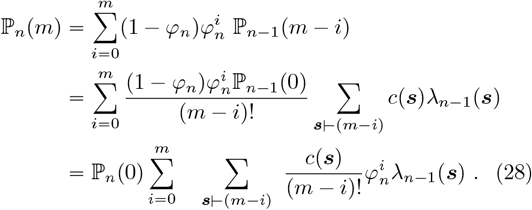

We shall show that this is in fact an alternative expression of Eq.(11).

From Eq.(22), we have the recurrence relationship

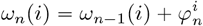

which by a standard expansion gives

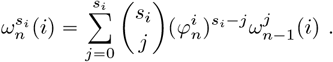

This immediately leads to

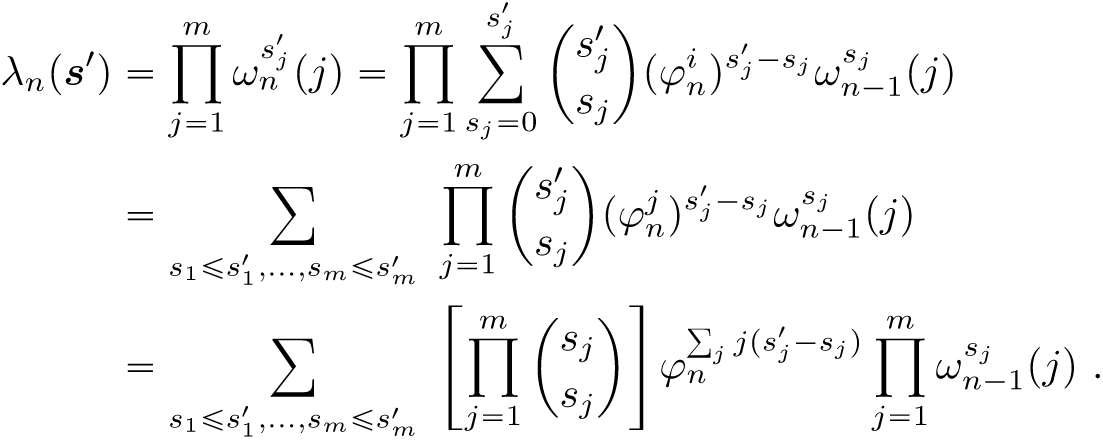

For two partitions 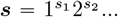 and 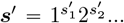,define the relationship ***s*** ⩽***s***^‵^ being true if and only if 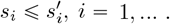, When ***s*** ⩽***s***^1^, define their difference partition as

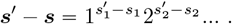

Consequently 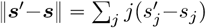,Since 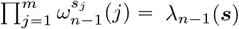, it follows that

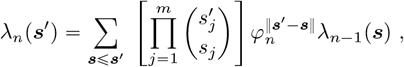

where the summation is taken over all partitions ***s*** ⩽***s***^‵^, which include the ∅. Grouping ***s*** according to its ‖ ***s*** ‖
results in

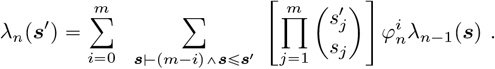

Therefore directly from (17) we have that

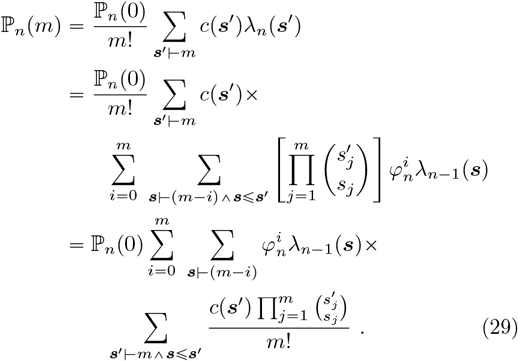

Since ***s*** ⊢ (*m* − *k*) and ***s***′ *m*, we have

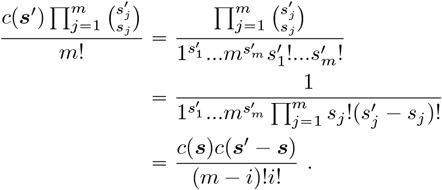

For a given ***s ⊢*** (*m* − *i*), enumerating over all ***s***^′^ ⊢ *m* with ***s***^′^ ⩾ ***s*** is equivalent to enumerating over all (***s***^′^ − ***s***) ⊢ *i*. Due to Eq.(20), it follows that

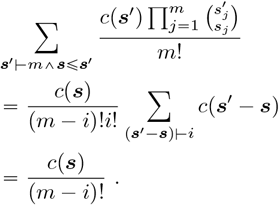

Therefore Eq.(29) is the same as Eq.(28), which indicates that Eq.(11) is also true for sample size *n*. By induction the Theorem is true for all sample sizes. ||

### Combinatorial interpretation

Mutations under the infinite-alleles model lead to partitions of the sample, resulting in the probability of a sample partition 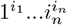 as given in Eq. (5), and the probability of having *k* distinct alleles as shown in Eq. (7). These probabilities have since been re-derived multiple times and subjected to various combinatorial interpretations (Hoppe, 1984; Donnelly, 1986; Donnelly and Tavaé, 1986; Griffiths and Lessard, 2005).

Under the infinite-alleles model, each distinct allele is unequivocally identified in the sample, but the number of mutations responsible for the observed allelic configuration is ambiguous—only the minimum number of mutations is known. In contrast, under the infinite-sites model, each mutation is unequivocally identified as a segregating site, but the number of mutations alone does not provide sufficient information on how the sequences in a sample are partitioned. This leads to an ambiguous sample configuration, unless more detailed information about each mutation is considered. Therefore, ℙ_*n*_(*m*) can only be partitioned according to the configurations of mutations. The theorem suggests that mutations can be partitioned into clusters corresponding to integer partitions, though the meaning of this is not entirely clear. Insight can be gained by examining in detail the cases of two and three mutations.

Consider the case of two mutations first. As for the states at which they may occur, there are only two possibilities. The first is that both mutations occur at the same state, and the second is that they occur at two different states. The probability that there are two mutations in the sample genealogy such that both occur at state *i* is

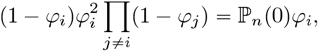

Therefore the probability that both mutations occur at the same state is ℙ_*n*_(0) ∑ *φ*_*i*_. Similarly the probability that there are two mutations in the sample genealogy such that one occurs at state *i* and another at state *j* is

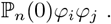

Enumerating over all *i* < *j* (since *j* < *i* and *i* < *j* represent the same event thus should not be counted twice), leads to that the probability two mutations occur at different state equal to 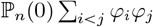.It follows that

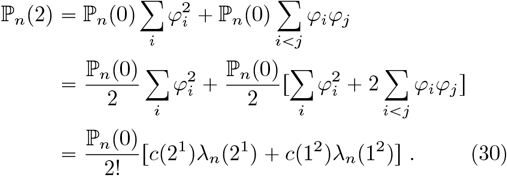

which is Eq.(11) when *m* = 2. The split of 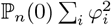 in the second equality into two halves, with the second half combined into the probability of two mutations at different states, is key to arriving at the concise expression in Eq. (30). One interpretation of this partition and reassembly is that the event of observing two mutations at the same state corresponds to the same outcome from two different underlying scenarios: one is that the two mutations arise in a tight cluster within a state, and the other is that the two mutations come from two separated clusters (each containing one mutation) at the same state. Since the sum of the probabilities of these two scenarios is 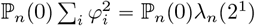,it is essential to understand how this probability is apportioned between the two scenarios. The derivation of Eq. (30) suggests an even split, which is equivalent to assigning the probability based on the proportion of the number *c*(2^1^) of permutations of cycle type 2^1^ among all permutations of two mutations, which is 2!. Therefore, if the two scenarios are separated, the scenario corresponding to a single cluster of two mutations has probability 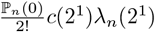 On the other hand, when the scenario of two separated clusters of one mutation each is combined with ℙ _*n*_(0) ∑ _*i*<*j*_ *φ*_*i*_*φ*_*j*_, it leads to a probability of 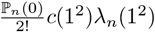,which represents the scenario where two mutations arise in two clusters, each containing one mutation. These clusters could either be at the same state or at different states.

The above analysis suggests that, although hypothetical and invisible, the concept of a cluster of mutations can help simplify the interpretation of mutations. The cluster of mutations should be the final assignment, as it will not be further divided or reassigned; otherwise, every mutation will eventually end up in its own cluster. Furthermore, each cluster resides within a single state. We can therefore refer to such clusters as atomic (indivisible) clusters. The rules for apportioning the probability of an event to different partitions of mutations into atomic clusters can be further clarified by examining the case of three mutations.

There are three different observable events for three mutations. The first is that all three occur at a single state, with probability 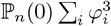,the second is that two mutations occur at one state and another at a different state, with probability 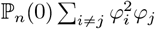,and the third is that three mutations occur at three different states, with probability ℙ _*n*_(0) ∑ _*i*<*j< k*_ *φ*_*i*_*φ*_*j*_*φ*_*k*_. The assembly of the probabilities leads to

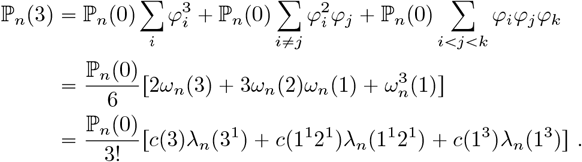

The first event is now split into three scenarios: one atomic cluster of three mutations, one atomic cluster of two mutations and one atomic cluster of one mutation, and three atomic clusters of one mutation each. The rule for appor-the total probability 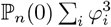 to the different scenarios is clearly proportional to the number of permutations of cycle types. Similarly, the second event is split into two scenarios: one atomic cluster of two mutations in one state and one cluster in a different state, and two atomic clusters of one mutation each in the same state and one cluster in a different state.

One can continue the analysis for more mutations, but the above analysis suffices the definition of **atomic clusters** as a list of mutations with the following three properties:

1. An atomic cluster cannot be further divided,
2. An atomic cluster resides within a single state,
3. The order (permutation) of mutations within an atomic cluster is cyclic.

A permutation is said to be cyclic if the elements are arranged in circular form so that it only matter what element follows a given element and there is no beginning nor end element. Therefore, a partition of *m* mutations into a group of atomic clusters corresponds to an integer partition 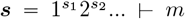.The number of permutations of *m* mutations to confirm with cycle type ***s*** can be intuitively derived as follows.

First of all, the number of arrangements of *m* mutations into *s*_*i*_ of groups (atomic clusters) of size *i*, (*i* = 1, …, *m*) is given by the standard multinomial coefficient

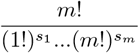

which assumes each group is labeled (identifiable). If groups can be identified only up to their sizes, then the total number of permutations is reduced to

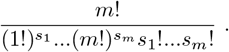

If permutation of mutations within each atomic cluster is identifiable, there will be more arrangements. The assumption of permutation within each atomic cluster being cyclic means that the number of permutations for atomic cluster of size *i* is (*i* − 1)!. Therefore the total number of arrangements of *m* mutations into cycle type ***s*** is

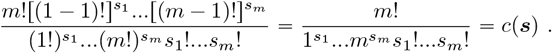

We thus arrive at a proposition that can be regarded as a corollary of the main theorem, based on the construct of hypothetical and invisible atomic clusters of mutations

#### Proposition.

*Mutations in the genealogy of a sample of n sequences emerge in atomic clusters. The specification of a list of atomic clusters coincides with an integer partition of the number of mutations m. The probability of mutations emerging in a partition* ***s*** *is given by*

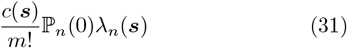

*and* ℙ _*n*_(*m*) *is the summation of probabilities of all different partitions of mutations, where c*(***s***), ℙ _*n*_(0) *and λ*_*n*_(***s***) *are defined, respectively, by (15), (12) and (16)*.

The construct of atomic clusters of mutations and Eq. (31) provide an analogy to Ewens’ sampling formula (5). Without the use of hypothetical and invisible boundaries to partition mutations into atomic clusters, attempting to express ℙ_*n*_(*m*) as the sum of probabilities for different scenarios of mutation realizations would lead to complicated, multi-layered summations, quickly losing simplicity.

### Numerical computation

When numerically evaluating ℙ _*n*_(*m*), it is important to verify its accuracy by comparing the results with known properties. In addition to the usual checks for a valid probability distribution, one can compare ℙ_*n*_(*m*) against its moments, including

1. **Expectation and variance** (Watterson, 1975; Fu, 2025):

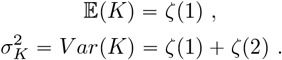
2. **Skewness and kurtonsis** (Fu, 2025):

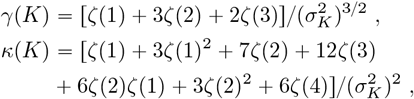

where

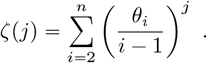

With these safeguards in place, we evaluated ℙ _*n*_(*m*) for various combinations of *n*,*m* and *θ*. In all cases, we encountered no issues with numerical accuracy when using Eq.(11). The impact of sample size *n* on the computation of ℙ _*n*_(*m*) is minor, as it only affects *ω*_*n*_(*i*), which needs to be evaluated once for each different *m*. However, the computational burden does increase with *m*, as will be discussed later.

We present two contrasting examples of ℙ _*n*_(*m*) here: one corresponding to the Wright-Fisher model with constant effective size, and the other to the constant-in-state model. Figure 1 shows the distribution of ℙ _*n*_(*m*) with *θ* = 3 for three different sample sizes: 50, 500, and 5000. These cases were chosen in part for comparison with the simulated data and the approximating normal distributions from Fu (2025) (Fig. 3, panel b).

**Fig. 1.**
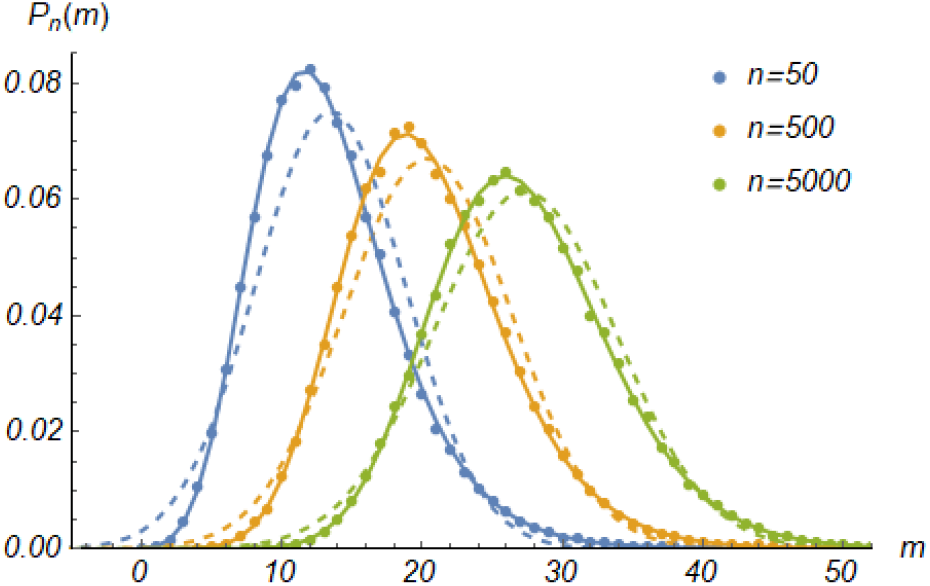
Probability distribution ℙ _*n*_(*m*) with *θ* = 3 for three samples sizes. For each sample size, solid curve is ℙ _*n*_(*m*) computed from (11), dot plot is from coalescent simulation and dashed curve represents the normal approximation. The latter two distributions are from panel (b) of Fig 3 in Fu (2025)

From Fig. 1, it is clear that the exact distribution ℙ _*n*_(*m*) for each sample size aligns well with the empirical distribution based on coalescent simulations of 50,000 replicates. This is expected, but without the exact distribution, one might wonder how many replicates are necessary to obtain sufficiently accurate results. With the exact distributions at hand, we can revisit the conclusions of Fu (2025) regarding the asymptotic normality of ℙ _*n*_(*m*). One conclusion in Fu (2025) is that the improvement towards normality from *n* = 50 to *n* = 500 is quite noticeable, while the further improvement from *n* = 500 to *n* = 5000 is more subtle. The patterns in Fig. 1 reinforce this conclusion with greater confidence. This is not surprising, given that the speed of c<u>onver</u>gence of ℙ _*n*_(*m*) to normality is approximately 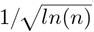. From to 500, the change is 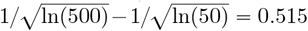,while from 500 to 5000 the change is only 0.426. Therefore, after an initial rapid improvement towards normality, further increases in sample size have progressively smaller effects.

For a population evolved according to the constant-in-state model, we considered three scenarios of population dynamics for *n* = 100. The first scenario represents a constant population size, with *θ*_*i*_ = 0.407 for *i* = 2, …, 15 and 2 for *i* = 16, …, 100. The second scenario corresponds to a recent population expansion,with *θ*_*i*_ = 0.407 for *i* = 2, …, 15 and 2 for *i* = 16, …, 100. The third scenario is a bottleneck followed by expansion, specified by *θ*_*i*_ = 0.65 for *i* = 2, 3, 0.1 for *i* = 4, 5, 6 and 1.425 for *i* > 6. All three scenarios have the same expected number of mutations *E*(*K*) = 5.177 The mean values of *θ* for these three scenarios are plotted in Fig. 2 against time into the past, scaled so that 1 unit corresponds to 4*N* generations, where *N* is the effective population size for the constant population. It is clear that, for the constant population size scenario, the average time to the most recent common ancestor (MRCA) is 1 unit (4*N* generations). In contrast, for scenarios 2 and 3, the time to the MRCA is shorter due to their smaller effective sizes near the MRCA.

**Fig. 2.**
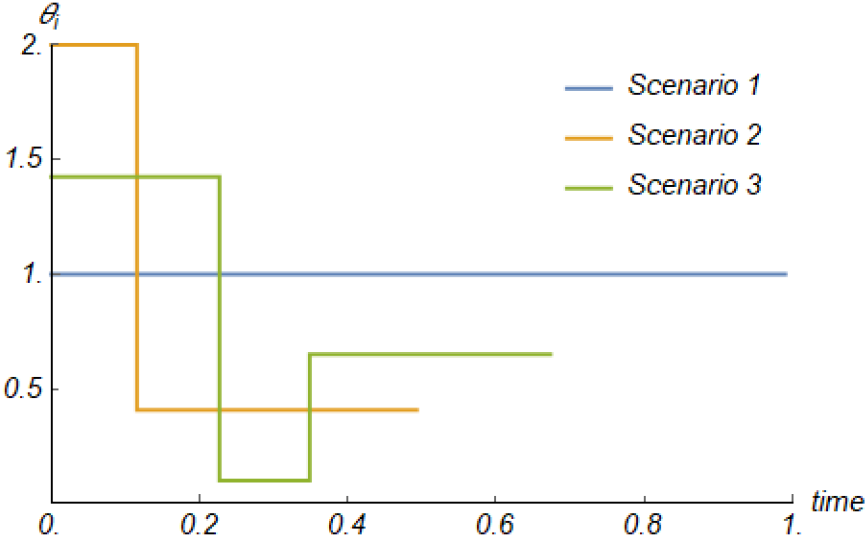
The mean dynamics of *θ*_*i*_ for a sample of size 100 for three scenarios. Scenario 1: constant model with *θ* = 1, Scenario 2: *θ*_*i*_ = 0.407, *i* = 2, …, 15 and *θ*_*i*_ = 2, *i* = 16, …, 100, and Scenario 3: *θ*_*i*_ = 0.65, *i* = 2, 3, *θ*_*i*_ = 0.1, *i* = 4, 5, 6 and *θ*_*i*_ = 1.425, *i* = 7, …, 100. All three scenarios lead to *E*(*K*) = 5.177.

Fig. 3 shows the distributions of ℙ _*n*_(*m*) for the three population dynamics scenarios described above. An immediate observation is that these distributions do not differ substantially, despite the seemingly dramatic differences in population size dynamics shown in Fig. 2. While there are noticeable differences around the peak of these distributions, the variations in the tails are minimal. This pattern serves as a reminder that *K* represents only one aspect of the genetic polymorphism within a sample. Behind *K*, the mutations constituting the sample can vary significantly in both type and frequency – a factor that has been leveraged in various data analysis approaches.

**Fig. 3.**
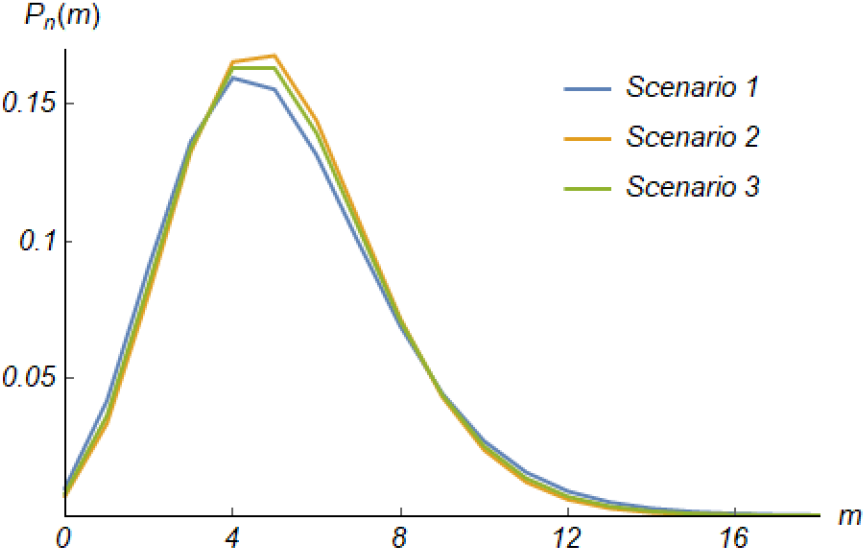
Probability distribution ℙ _*n*_(*m*) computed from (11) for n=100 and three scenarios in Fig. 3

#### Impact of integer partitions

A key technical issue affecting the computation of ℙ _*n*_(*m*) is the number and patterns of integer partitions. Let *p*(*m*) denote the number of different partitions of the integer *m*. The value of *p*(*m*) is a critical factor in determining the computational complexity of ℙ _*n*_(*m*) using Equation (11), as the computation requires enumerating all possible partitions of *m*. The function *p*(*m*) is a classic problem in combinatorics and number theory, and while a closed-form formula for *p*(*m*) is not available, it can be easily computed through enumeration or recurrence relations. For example, the only partition for *m* = 0 is the empty set ∅, *p*(1) = 1 corresponds to the partition (1), and *p*(2) = 2 corresponds to the partitions (2) and (1, 1). While *p*(*m*) increases slowly for small values of *m*, it grows rapidly as *m* increases. The thresholds of 10^3^, 10^6^ and 10^9^ partitions are reached at *m* = 22, 61, and 115, respectively. Table 1 provides values of *p*(*m*) for selected values of *m* up to 150 (Sloane, 2024; Wolfram Research, 2014).

**Table 1.** The number *p*(*m*) of integer partitions for selected *m* ⩽150.

| m | p(m) | m | p(m) | m | p(m) |
| --- | --- | --- | --- | --- | --- |
| 0 | 1 | 15 | 175 | 30 | 5,604 |
| 1 | 1 | 16 | 231 | 35 | 14,883 |
| 2 | 2 | 17 | 297 | 40 | 37,338 |
| 3 | 3 | 18 | 385 | 45 | 89,135 |
| 4 | 5 | 19 | 490 | 50 | 204,226 |
| 5 | 7 | 20 | 627 | 55 | 451,276 |
| 6 | 11 | 21 | 792 | 60 | 966,467 |
| 7 | 15 | 22 | 1,002 | 65 | 2,012,558 |
| 8 | 22 | 23 | 1,255 | 70 | 4,087,968 |
| 9 | 30 | 24 | 1,575 | 75 | 8,118,264 |
| 10 | 42 | 25 | 1,958 | 80 | 15,796,476 |
| 11 | 56 | 26 | 2,436 | 90 | 56,634,173 |
| 12 | 77 | 27 | 3,010 | 100 | 190,569,292 |
| 13 | 101 | 28 | 3,718 | 125 | 3,163,127,352 |
| 14 | 135 | 29 | 4,565 | 150 | 40,853,235,313 |

Since every partition contributes positively to the overall probability ℙ_*n*_(*m*), there is generally no issue with error propagation when using double-precision computation in modern programming languages and tools. However, for *m* > 50, *p*(*m*) increases rapidly, presenting a significant computational challenge. For instance, when *m* = 60, the number of partitions is nearly 1 million, and by the time *m* reaches 100, the number swells to 190 million. To make matters even more daunting, for *m* = 150, there are over 40 billion partitions! While large numbers may offer elegant solutions in certain contexts, such as in the application of the central limit theorem, they become a major hurdle in the numerical computation of ℙ_*n*_(*m*). Therefore, investigating the roles of different types of partitions in the computation of ℙ_*n*_(*m*) is important for understanding and improving the efficiency of these calculations.

Two important characteristics of integer partitions are partition length, which refers to the number of parts in the partition, and maximal size, which denotes the largest value of the parts. In the context of mutation partitions, these correspond to the number of atomic clusters and the maximum number of mutations in a single atomic cluster, respectively. Let *p*_*i*_(*m*) represent the number of partitions of length *i*, and 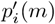 denote the number of partitions with a maximal size of *i*. It is known that 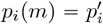 (*m*) (Slomson, 1991), and thus

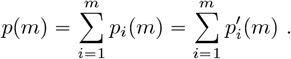

Then *p*_*i*_(*m*)/*p*(*m*) is the percentage of partitions of length *i* among a total *p*(*m*) partitions. Similarly 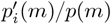 is the percentage of partitions with maximal size *i* among *p*(*m*). For a partition 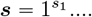,define #(***s***) as its number of parts and #^‵^ (***s***) as its maximal size of parts. Then

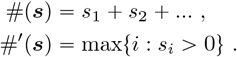

Define ℙ_*n*_(*m, i*) be the contribution to ℙ_*n*_(*m*) by partitions of length *i*, and similarly P^1^ (*m, i*) be the contribution to ℙ_*n*_(*m*) by partitions ***s*** with maximal size *i*. Then

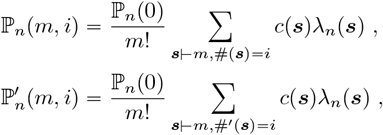

and 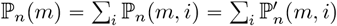

Plots of the *p*_*i*_(*m*)/*p*(*m*) against *i*/*m* for three different *m* with *n* = 500 and *θ* = 10 are given in Fig. 4. It is clear that partitions of length in the range 0.2*m* −0.3*m* are most abundant. Take the case of *m* = 100 for example (green dotted plot), the peak of *p*_*i*_(*m*)/*p*(*m*) is around 0.2 for *i*/*m*, which means the highest concentration of partitions are those of length 20. Additionally, it is apparent that as *m* increases, the peak concentration of partitions becomes smaller. Fig. 4 also plots ℙ_*n*_(*m, i*)/ ℙ_*n*_(*m*) against *i*/*m*. The contribution towards ℙ_*n*_(*m*) is concentrated around 0.55 in the case of *m* = 100 (green curve), indicating that partitions of length around 55 contribute most. It is also clear that the peak of partitions contributing to ℙ_*n*_(*m*) gradually shifts to the left with increase of *m*. Furthermore the density *p*_*i*_(*n*)/*p*(*m*) and the density ℙ_*n*_(*m, i*)/ ℙ_*n*_(*m*) for a given *m* overlap only slightly, meaning that in general a small fraction of integer partitions contribute to the overwhelming portion of ℙ_*n*_(*m*). In other words, ℙ_*n*_(*m*) can typically be computed with sufficient accuracy by evaluating a small portions of partitions. For example, for *m* = 100, 99.99% of the ℙ_500_(100) comes from the partitions of length 30 or larger, which account for only 13.4% of the total 190 millions partitions of 100. This feature is quite beneficial, as it makes the challenge of dealing with a large number of partitions a manageable issue.

**Fig. 4.**
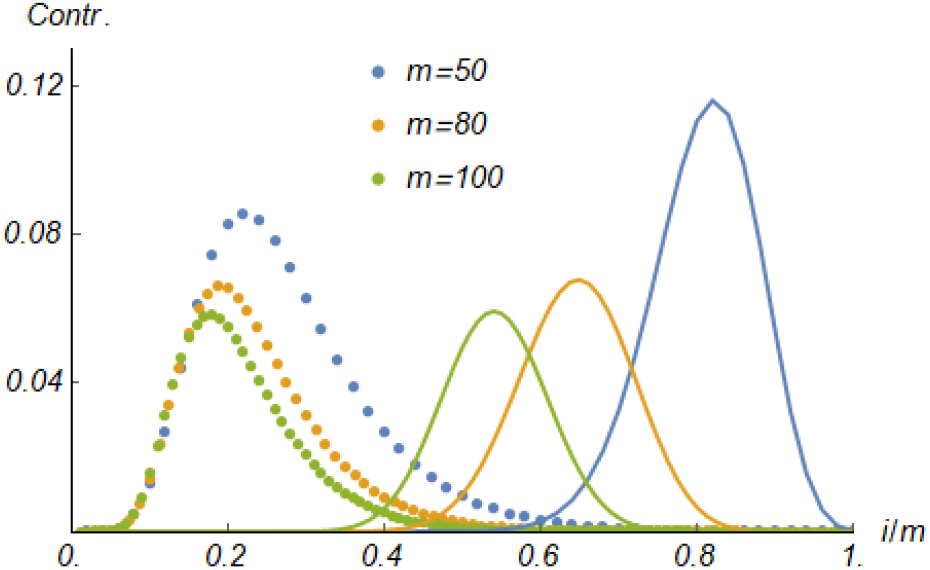
Percentage of partitions of different lengths (dots) and their contributions(lines) to ℙ_*n*_(*m*), for *n* = 500 and *θ* = 10. Three numbers of mutations considered are *m* = 50, 80 and 100.

To test the limits of evaluating ℙ_*n*_(*m*) using Eq. (11), we computed ℙ_500_(150), which required enumerating over 40 billion partitions of 150. This case was used to examine both the contribution of partitions of different maximal sizes to ℙ_*n*_(*m*) and the mean contribution of partitions of various lengths. The results are presented in Fig. 5. From the figure, it is evident that the contribution by partitions of different maximal sizes (green curve) closely overlaps with the percentage of partitions of various maximal sizes (orange curve). This suggests that prioritizing the enumeration of partitions based on their maximal size does not lead to a significant reduction in computational effort.

**Fig. 5.**
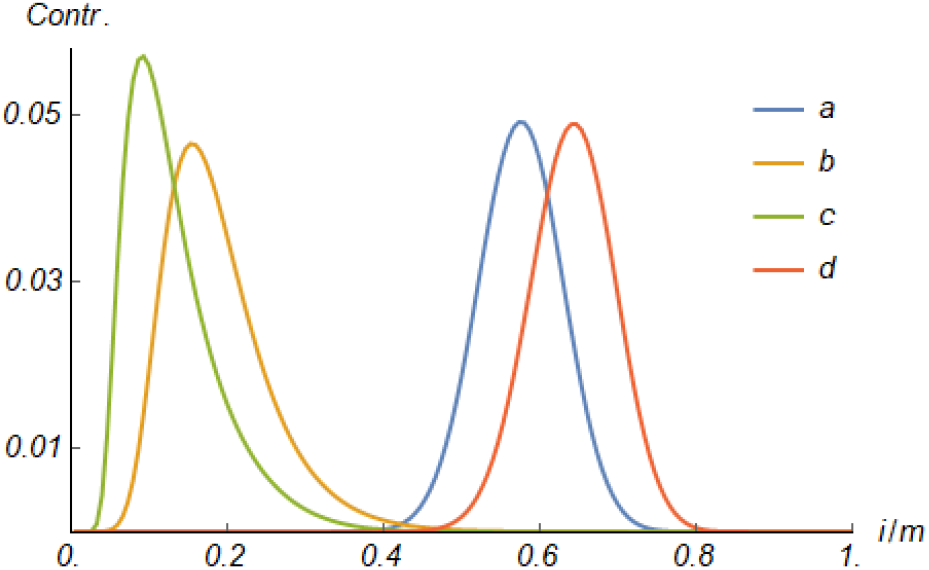
Contributions of partitions to ℙ_500_(150) = 0.01044 with *θ* = 20. (a) Contribution of partitions of different lengths, (b) percentage of *p*_*i*_(*m*) in *p*(*m*), (c) Contribution by partitions of different maximal sizes and (d) Contribution per partition of different length.

Fig. 5 also shows the per-partition contribution to ℙ_*n*_(*m*) for different lengths (red curve). Interestingly, this curve does not coincide with the overall contribution of different lengths (blue curve). In other words, the partition lengths that contribute most to ℙ_*n*_(*m*) are not the same as those that contribute the most per partition. However, it is clear that, whether considered collectively or individually, partitions with lengths equal to or larger than *m*/2 contribute most to ℙ_*n*_(*m*).

#### Conditions for the failure of Tavaré formula

It may seem straightforward to compute ℙ_*n*_(*m*) using Eq.(4) under the Wright-Fisher model with constant effective size, which is the sum of *n* − 1 summands. However, once sample size *n* is sufficiently large, the binomial coefficient 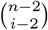 in the (*i* − 1)-th summand can be a very large number, particularly when *i* is around n/2. As a result, Tavaré formula can often be a summation of some huge summands with alternating sign. To make the matter worse, adjacent summands of opposite signs are typically not close enough in value to cancel each other quickly. When ℙ_*n*_(*m*) is about a magnitude 10^−15^ smaller than the absolute values of some summands, error accumulation/propagation is inevitable. This is because modern computers generally carry out floating point calculation in 64-bits format, which leads to 15-17 significant digits, regardless of program languages/tools. There have been improvements in various programming languages, but they are mostly on integer operations and data format. Once floating point operations are performed, the notorious error issue persists.

For illustration, consider ℙ_100_(5) with *θ* = 1. The 4 summands of Eq.(4) for i=48 to 51 are, respectively, − 8.31062×10^17^, 7.66893×10^17^, −6.81003×10^17^, 5.81884× 10^17^. When adding summands in ascending order of *i*, the result is 7.089 ×10^4^ in standard double precision Java programming, which leads to ℙ_*n*_(5) = 7.01 × 10^6^, while the correct value is ℙ_100_(5) = 0.1553 according to Eq. (11). One can explore different orders of adding summands and the final result does change slightly, but all results in qualitatively wrong answer.

In our attempt to improve the applicability of Eq.(4), we explored three different implementations, which were standard programming in Java (version 22) using double precision, Java programming using BigDecimal class which has unlimited precision for integer computation and improved precision for decimal numbers, and symbolic programming by Mathematica V10 (Wolfram Research, 2014), which is supposed to have unlimited precision in applicable numerical computations. We observed that significant error usually started with smaller *m*, as expected, because the power factor 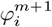 with large *m* can help to lower the absolute value of a summand, thus reducing the error accumulation. Also expected is that for a given combination of *m* and *θ*, errors become more pronounced with increasing sample size. This prompted us to exam the threshold sample size that leads to significant error in ℙ_*n*_(*m*). Table 2 gives the these sample sizes for several values of *θ* in ℙ_*n*_(1) and ℙ_*n*_(5). We started with *m* = 1 instead of *m* = 0 since ℙ_*n*_(0) can be computed directly by Eq.(3) without using Eq.(4). We define failure of Eq.(4) as a situation in which ℙ_*n*_(*m*) has at least 10% error when compared with ℙ_*n*_(*m*) from Eq.(11).

**Table 2.** Failure conditions for computing ℙ_*n*_(*m*) using Eq.(4) by three different programming methods.

| $\theta$ | Java double | BigDecimal | Mathematica |
| --- | --- | --- | --- |
| 0.05 | 57<sup>a</sup>,80<sup>b</sup> | 60,87 | 62,82 |
| 0.10 | 58,80 | 59,83 | 58,83 |
| 0.50 | 52,75 | 58,80 | 58,81 |
| 1.00 | 48,70 | 55,75 | 54,76 |
| 5.00 | 38,52 | 39,55 | 40,55 |
| 10.0 | 27,38 | 30,45 | 30,41 |

Note. Error rate = $100 \times [\mathbb{P}'_n(m) - \mathbb{P}_n(m)] / \mathbb{P}_n(m)$ where $\mathbb{P}_n(m)$ is computed by Eq.(11) and $\mathbb{P}'_n(m)$ by Eq.(4). <sup>a</sup> sample size at or larger will lead to $\mathbb{P}'_n(1)$ with at least 10% error, <sup>b</sup> sample size at or larger will lead to $\mathbb{P}'_n(5)$ with at least 10% error.

It is clear from Table 2 that for a wide range of *θ*, Eq.(4) starts to fail for ℙ_*n*_(1) for sample size around 30 − 60, and starts to fail for ℙ_*n*_(5) for sample sizes around 40 − 85. Higher precision programmings push the threshold sample sizes only slightly higher, which validates the early statement that internal computation in modern computers is mostly 64-bits format, regardless of programming languages/environments. Consequently Tavaré formula is not recommended in general for numerical computation when *n* > 50, particularly when the entire range of *m* need to be considered.

## Summary

A new formula is derived in this paper for the probability of the number *K* of mutations in the genealogy of a random sample of DNA sequences from a single locus without recombination, drawn from a population evolved according to the constant-in-state model. In the case of constant effective population size, this formula closely resembles Ewens’ sampling formula for the number of distinct alleles in the sample. The new sampling formula is based on the probabilities of different partitions of mutations into coalescent states and achieves significant simplification by recognizing and partitioning mutations into atomic clusters – which are indivisible, bounded within coalescent states, and where permutation of mutations is cyclic. Without the concept of atomic clusters, expressing ℙ_*n*_(*m*) as the sum of probabilities for different realizations of mutations in the sample genealogy would lead to complex, multi-layered summations whose complexity grows rapidly with the number of mutations.

The new formula can be used effectively for numerical computation with minimal concern for accuracy, since every summand contributes positively. While the efficiency of the computation remains largely unaffected by sample size, it is impacted by the number of mutations due to the need to enumerate all possible partitions of the given number of mutations. In typical applications, the formula should handle computations efficiently with little or no optimization of the algorithm. Moreover, partitions with a larger number of atomic clusters contribute more to the overall probability, so computation efficiency can be improved by prioritizing the evaluation of partitions based on the number of atomic clusters.

In contrast, Tavaré’s formula has a limited scope of applicability, primarily due to the error-prone summation of large summands with alternating signs. As such, it is not recommended for general numerical computation of the distribution of *K*, especially when the sample size is 50 or greater. However, it may still be useful for computing ℙ_*n*_(*m*) for large *m* with smaller sample sizes, potentially in conjunction with the new formula. A Java package implementing the new formula for ℙ_*n*_(*m*) is available from the author upon request.

The relevance of permutations of cycle type and integer partitions to population genetics was first highlighted through Ewens’ sampling formulas under the infinite-alleles model (Ewens, 1972; Karlin and McGregor, 1972), but has since had limited application to other summary statistics. The newly derived sampling formula also successfully connects to permutations of cycle type and integer partitions. The links between theoretical population genetics and the fields of combinatorics and number theory may be deeper than previously recognized. Consequently more summary statistics related to *K* can be expected to possess such characteristics.

## Notes

### Competing Interest Statement

The authors have declared no competing interest.

### Summary of Updates

A few typos in the first two sections were corrected. Although they do not alter the flow of the manuscript in meaningful way, the correction should help to improve readability of the manuscript.

